# PqsE is conserved and functionally relevant in *Pseudomonas aeruginosa*

**DOI:** 10.1101/2025.05.16.654539

**Authors:** Mylène C. Trottier, Marie-Christine Groleau, Jeff Gauthier, Antony T. Vincent, Roger C. Levesque, Eric Déziel

## Abstract

The bacterium *Pseudomonas aeruginosa* is an opportunistic pathogen responsible for several acute and chronic infections. It produces a diverse array of virulence and survival determinants, many of which are tightly regulated by three interlinked quorum sensing (QS) systems named *las, rhl* and *pqs*. RhlR, the transcriptional regulator of the *rhl* system, activates multiple virulence genes upon binding to its cognate autoinducer signal. Meanwhile, the *pqs* system relies on 4-hydroxy-2-alkylquinolines (HAQs) as signaling molecules to induce the transcriptional regulator MvfR (PqsR). MvfR then activates the transcription of the *pqsABCDE* operon, encoding enzymes for HAQ biosynthesis. The final gene in this operon encodes PqsE, a multifunctional protein unique to *P. aeruginosa*. Beyond its thioesterase activity in HAQ biosynthesis, PqsE stabilizes RhlR, facilitating its regulation of target genes, some of which are implicated in virulence. Because of its role in pathogenicity, PqsE is regarded as a promising therapeutic target for combating *P. aeruginosa* infections.

While the role of PqsE towards the RhlR regulon is increasingly understood in prototypical *P. aeruginosa* strains such as PA14 and PAO1, its broader relevance as an anti-virulence target remains underexplored. Here, we confirm that PqsE is functionally relevant across a panel of twelve genetically diverse *P. aeruginosa* strains using metabolite quantification and phenotypic assays. Significant strain-to-strain variations further highlight the importance of studying QS regulation among diverse isolates. Moreover, this study underscores PqsE as a key QS regulator with a conserved role in coordinating virulence determinant production and social behaviors across diverse *P. aeruginosa* populations.

**IMPORTANCE:** *Pseudomonas aeruginosa* is a versatile opportunistic pathogen, naturally tolerant and readily acquiring resistance to multiple antibiotics. Consequently, the World Health Organization identified this bacterium as a high priority pathogen for researching and developing new antimicrobial strategies. *P. aeruginosa* utilizes quorum sensing, a cell-to-cell communication system, to regulate the expression of several of its virulence factors. Here, we confirm that the PqsE protein is conserved and that its function in quorum sensing, especially towards the RhlR regulator, is maintained across a panel of twelve genetically diverse *P. aeruginosa* strains. Since PqsE is conserved and unique to this bacterium, it could serve as an ideal target for anti-virulence therapies, offering new alternatives to combat antimicrobial resistance.

## INTRODUCTION

The bacterium *Pseudomonas aeruginosa* is an opportunistic pathogen closely associated with human activity and is a leading cause of infections, both acute and chronic, in immunocompromised individuals (1, 2). Its capacity to adapt to various environments and produce virulence factors is mainly regulated by a process named quorum sensing (QS), a cell-to-cell communication system that modulates gene expression through the production and detection of small autoinducer molecules in response to population density (3, 4).

*P. aeruginosa* has three interdependent QS systems: *las*, *rhl*, and *pqs*. The *las* system relies on the LasI synthase, which produces the signal molecule 3-oxo-dodecanoyl-homoserine lactone (3-oxo-C_12_-HSL). This molecule binds and activates LasR, its cognate LuxR-type transcriptional regulator, which in turn promotes the expression of numerous genes, including *lasI*, thereby sustaining autoinducer production through a positive feedback loop (5, 6). In prototypical strains, LasR activates the *rhl* system by initiating the transcription of *rhlI* and *rhlR* (7, 8). The RhlI synthase produces the butanoyl-homoserine lactone (C_4_-HSL) autoinducer, while RhlR functions as a transcriptional regulator. Upon binding to C_4_-HSL, RhlR induces the expression of *rhlI* and genes responsible for the production of virulence factors, including those involved in pyocyanin synthesis (two *phzABCDEFG* operons) and the production of rhamnolipids (*rhlAB*, *rhlC*) (6, 8–11). While the *las* system is generally considered atop the QS hierarchy in prototypical strains, loss of LasR activity is frequent in strains isolated from both clinical and environmental settings (12–15). Nevertheless, some LasR-defective strains still retain RhlR function and fully express virulence factors, suggesting that RhlR could assume a central role in a malleable QS hierarchy (13–15). The third QS system in *P. aeruginosa,* the *pqs* system, is driven by signaling molecules called 4-hydroxy-2-alkylquinolines (HAQs). The transcriptional regulator of this system, MvfR (also known as PqsR), is activated upon binding either 4-hydroxy-2-heptylquinoline (HHQ) or 3,4-dihydroxy-2-heptylquinoline (*Pseudomonas* Quinolone Signal; PQS) (16). Once bound to one of its autoinducing ligands, MvfR regulates the transcription of the *pqsABCDE* operon, which encodes the enzymes responsible for HAQ biosynthesis (17–20). Furthermore, the *pqs* system is thoroughly regulated by the two other QS systems, as LasR positively regulates both the transcription of *mvfR* and the *pqsABCDE* operon, whereas RhlR negatively regulates the expression of *pqsABCDE* (20–22).

The final gene in the *pqsABCDE* operon, *pqsE*, encodes the multifunctional protein PqsE. It was characterized as a thioesterase implicated in HAQ biosynthesis (23). However, PqsE seems dispensable for this process, as *pqsE* mutants show no defects in HAQ production in prototypical strains, presumably because redundant enzymes can take over its function (18, 19, 23). In addition to its enzymatic activity, PqsE modulates the transcription of multiple target genes, most of which belong to the RhlR regulon, including the *phzABCDEFG* operons, and to some extent, the *rhlAB* operon (19, 24–29). Indeed, PqsE has a minimal impact on the *P. aeruginosa* transcriptome in the absence of RhlR (24). Recent studies have revealed that PqsE interacts directly with RhlR through a protein-protein interaction, enhancing RhlR stability and increasing its affinity for target promoters (26, 30–32). The chaperone-like activity of PqsE is particularly noteworthy, as it is unique to *P. aeruginosa* (30). Notably, the PqsE-RhlR complex functions independently of the catalytic thioesterase activity of PqsE (30, 31, 33). Furthermore, an implication for PqsE in biofilm formation and virulence has also been reported in strains PA14 and PAO1 (25, 28, 34, 35).

Due to its specificity and role in virulence, PqsE is often considered an attractive target for the development of anti-virulence therapies (26, 30, 31, 36, 37). However, although the functional role of PqsE towards the RhlR regulon and HAQ production has been extensively studied in prototypical *P. aeruginosa* strains PA14 and PAO1, its role in other genetically diverse isolates has not been explored. To establish its potential as a target for anti-virulence strategies, it is essential to first confirm the genetic and functional conservation of PqsE across diverse strains.

In this study, we conducted a genetic and functional analysis of PqsE in diverse isolates to gain knowledge on its ecological role. We focus on virulence determinants reported as being regulated by PqsE in prototypical strains. Our results show that the *pqsE* gene is conserved, but the functional role of PqsE generally varies across isolates, although its importance in regulating some virulence determinants and social behaviours is widespread.

## RESULTS AND DISCUSSION

### The pqsE and rhlR genes are conserved in genetically different P. aeruginosa backgrounds

To be considered a key target for developing anti-virulence strategies, PqsE would need to be genetically and functionally conserved. Thus, a panel of twelve *P. aeruginosa* strains, including PA14 and PAO1, was selected to study the functionality and conservation of PqsE (**Table 1; Table S1)**. These strains were chosen to capture ecological and genetic diversity and were obtained from various clinical and environmental contexts. Some of them also display diverse QS functional patterns. Indeed, our panel includes LasR-defective strains, with or without an independently active RhlR, and one strain unable to produce HAQs, as determined previously (13, 14). Notably, whole-genome sequences are available for all strains (38).

**Table 1.** Characteristics of P. aeruginosa isolates used in this study.

| Strains | Origin | Pathology or environment | LasR classification* | Production of HAQs* | Source |
| --- | --- | --- | --- | --- | --- |
| **Prototypical strains** |  |  |  |  |  |
| PA14 | USA | Burn wound | Functional LasR | Yes | (39) |
| PAO1-N (PAO1) | Australia | Wound infection | Functional LasR | Yes | G. Rampioni |
| **Clinical strains** |  |  |  |  |  |
| 39016 | United Kingdom | Keratitis | Functional LasR | Yes | (38) |
| A22 | France | Wound | Functional LasR | No | (40) |
| E90 | USA | Cystic fibrosis | LasR-defective, RhIR-active (RAIL) | Yes | (15) |
| JJ692 | USA | Urinary tract infection | LasR-defective | Yes | (41) |
| PA-W9 | United Kingdom | Leg Ulcer | Functional LasR | Yes | (38) |
| SMC1596 | Canada | Cystic fibrosis | Functional LasR | Yes | (42) |
| **Environmental strains** |  |  |  |  |  |
| PA-CL512 | Canada | Hospital sink | LasR-defective, RhIR-active (RAIL) | Yes | (43) |
| PA-CL513 | Canada | Hospital sink | Functional LasR | Yes | (43) |
| PA-CL521b | Canada | Hospital sink | LasR-defective, RhIR-active (RAIL) | Yes | (43) |
| PG201 (Rsan-ver) | Switzerland | Soil | LasR-defective | Yes | (44) |

\*Classification of LasR function and HAQ production is based on Groleau et al. (2022) (13) and Trottier et al. (2024) (14).

To gain deeper insight into the evolutionary dynamics of this panel of strains, we conducted a phylogenetic analysis based on the conservation of the core genome. The generated distance tree provides information about the core genome diversity of the analyzed *P. aeruginosa* strains (**Fig. 1)**. Many strains exhibit a high core genome similarity, suggesting limited divergence in conserved functional pathways. However, strains located on distinct branches of the tree (e.g., the PA14 group vs. the PAO1 group) may possess slight differences in their set of genes, explained by evolutionary divergences.

**Figure 1.**
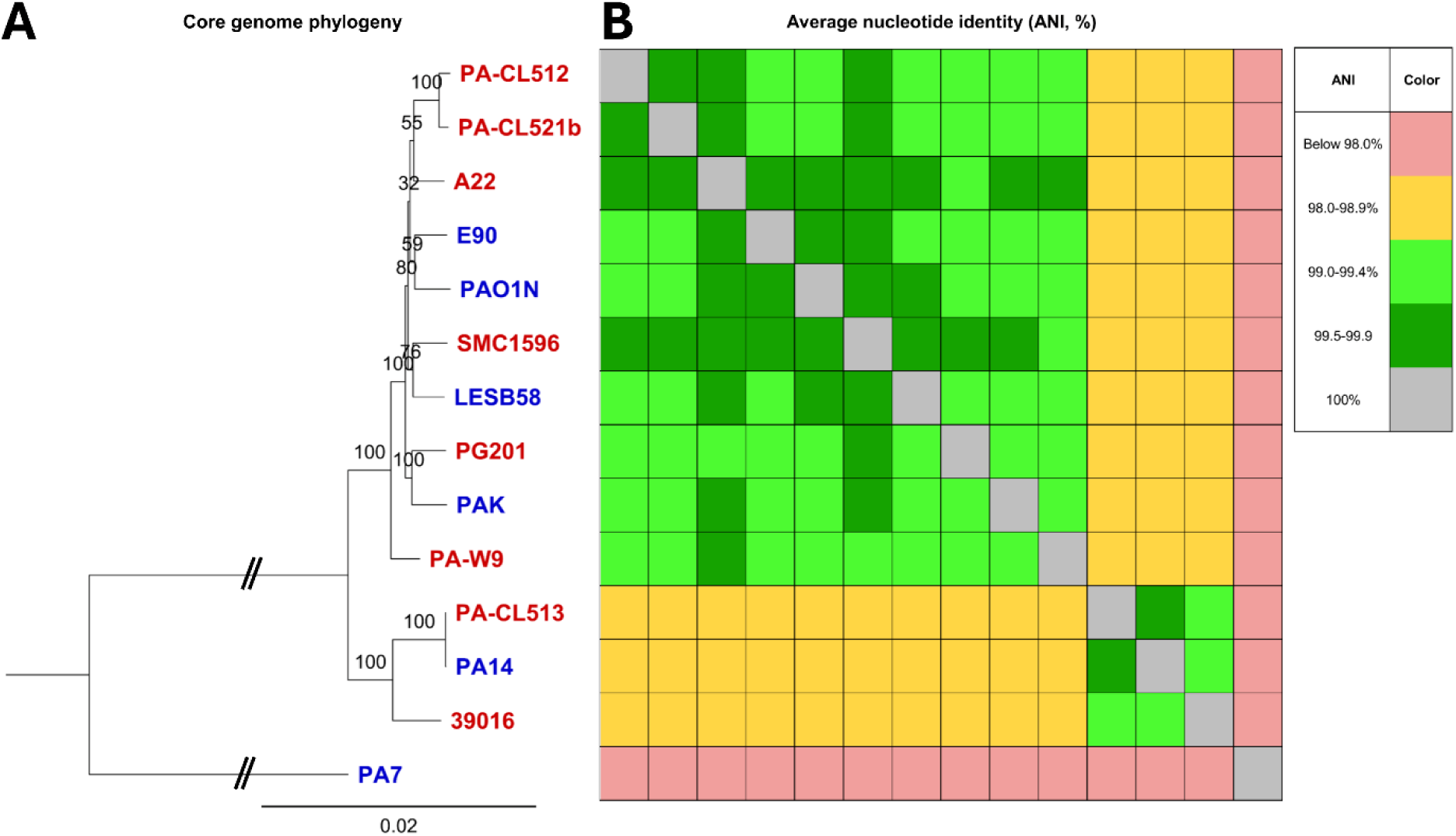
**Core genome phylogeny and ANI heatmap of selected *P. aeruginosa* strains. A: Core genome phylogenetic tree of *P. aeruginosa* strains from this study (red), along with prototypical strains (blue)**. Node labels indicate statistical support (in %) among 1,000 bootstrap replicates. Core genes included in this phylogeny were inferred from a core gene alignment generated by PyMLST v2.1.6 using the cgMLST.org public scheme for *P. aeruginosa,* which includes 3,687 core loci from 6,730 genomes. The GTR+F+I+G4 model was selected according to the Bayesian Information Criterion by IQ-TREE v2.3.6 as the best-fit model. Branch lengths indicate the number of substitutions per site. **B: Average nucleotide identity matrix aligned with the phylogenomic tree in A.** ANI calculations were done with pyani v2.1.6 using the “ANIm” all-versus-all comparison mode, with default parameters.

None of our strains cluster with the PA7 outlier group (recently renamed as the *Pseudomonas paraeruginosa* group (45)). Within the main branch, the strains form two distinct clusters: one that includes 10 clinical and environmental strains along with PAO1, and another where two strains cluster with PA14, thus confirming that all strains used in this study accurately belong to the *P. aeruginosa* species. As expected, environmental and clinical strains do not cluster separately (38). Also, patterns of QS function (LasR-defective, LasR-defective RhlR-active (RAIL), LasR-positive) generally do not correlate with the phylogenetic clustering. However, one distinct cluster composed of PA-CL513, PA14 and 39016 stands out, as all three strains are LasR-positive and produce HAQs. These findings further support that there are some variations within the core genomes of our strains, which could further explain phenotypic differences among them. These potential differences are also reflected in the broad average nucleotide identity range both within and between each *P. aeruginosa* core phylogenomic cluster (99.1 to 99.7%) (**Fig. 1B).**

Since previous studies have shown that *pqsE* is highly conserved across diverse *P. aeruginosa* isolates (46, 47), we aimed to confirm whether conserved mutations or polymorphisms were present. To do so, we analyzed the genomic sequence of *pqsE* and the predicted amino acid sequence of the PqsE protein across our panel of *P. aeruginosa* strains. Our genomic analysis confirms that *pqsE* is constitutively present and highly conserved, revealing a 99.5% (902/906 nucleotides) identity across strains. Three nucleotide positions with polymorphisms among the strains and one unique mutation were identified **(Table S2)**. The identified polymorphisms do not affect the predicted amino acid sequence **(Fig. S1)** or codons encoding amino acids crucial for interactions with RhlR, as identified in prior studies (26, 30, 32). The only exception is strain SMC1596, harboring a SNP at position 516, located within a codon critical for residue R172 of the PqsE protein, a position important for interacting with RhlR (30). However, because this SNP is non-synonymous, there is no impact on the predicted amino acid sequence, confirming the functional conservation of this interaction site across isolates **(Fig. S1, Table S2)**.

Since PqsE and RhlR function as a pair to regulate the transcription of various genes, and variations in *rhlR* could thus impact PqsE-dependent phenotypes, we also verified the conservation of the *rhlR* gene. Indeed, this gene has a low mutation rate according to the literature (48, 49). Like *pqsE*, our analysis revealed only few polymorphisms in *rhlR* sequence **(Table S3)**, that have no impact on the predicted amino acid sequence **(Fig. S2)**.

Overall, our results reveal that the predicted PqsE and RhlR proteins are conserved in our panel of diverse strains. Importantly, previously identified codons encoding key amino acids involved in PqsE-RhlR interactions are preserved **(Fig. S1, Fig. S2)** (30). From a strictly genetic perspective, *pqsE* remains a promising target for anti-virulence therapies due to its high conservation, as shown here and in prior studies, and its specificity to *P. aeruginosa* (46, 50).

### PqsE impacts the production of virulence determinants in P. aeruginosa isolates

While *pqsE* is genetically conserved, the functional conservation of its corresponding protein, PqsE, remains unclear. We first examined the production of virulence determinants regulated by PqsE in prototypical strains, such as pyocyanin and rhamnolipids. This approach allowed us to determine whether phenotypes observed in a couple of strains could be reproduced in genetically diverse isolates. We included PA14 and PAO1 in our panel of strains, since their PqsE-dependent phenotypes are well described (19, 25, 27–29). To investigate the role of PqsE, we deleted *pqsE* in all strains from our panel. Importantly, we confirmed that Δ*pqsE* mutants exhibited no differences in growth compared to their respective wild-type (WT) counterparts **(Fig. S3)**. We also included the previously characterised Δ*pqsE* mutants of PA14 and PAO1 (18, 25). The next step was to phenotypically characterize all the generated Δ*pqsE* mutants and verify if the impact on regulated traits was conserved between isolates.

The most studied PqsE-dependent phenotype is pyocyanin production. Pyocyanin is a redox-active phenazine responsible for the characteristic blue pigmentation of *P. aeruginosa* cultures. Its biosynthesis involves two redundant operons, *phzA1-G1* (*phz1*) and *phzA2-G2* (*phz2*), which code for enzymes to produce phenazine-1-carboxylic acid (PCA), a precursor that is subsequently converted to pyocyanin by PhzM and PhzS (51). The *phz1* operon possesses a *lux*-box recognized by RhlR and is the predominant *phz* operon expressed in planktonic cultures of the PA14 strain (52, 53). Regulation of the expression of *phz1* in prototypical strains depends on the functionality of PqsE on RhlR activity (29, 54, 55).

To investigate the role of PqsE in this process across our panel of strains, we quantified PCA, the direct product of the *phz1* operon, along with its final metabolite, pyocyanin, using liquid chromatography coupled to tandem mass spectrometry (LC/MS/MS) in WT strains and their isogenic Δ*pqsE* mutants. In 11 out of the 12 strains, we measured a significant reduction or complete loss of PCA and pyocyanin in the Δ*pqsE* mutant, at least for one of the two time points (**Fig. 2A)**. This widespread reduction confirms that the role of PqsE in promoting pyocyanin biosynthesis is broadly conserved across diverse *P. aeruginosa* isolates, consistent with previous findings (29).

**Figure 2.**
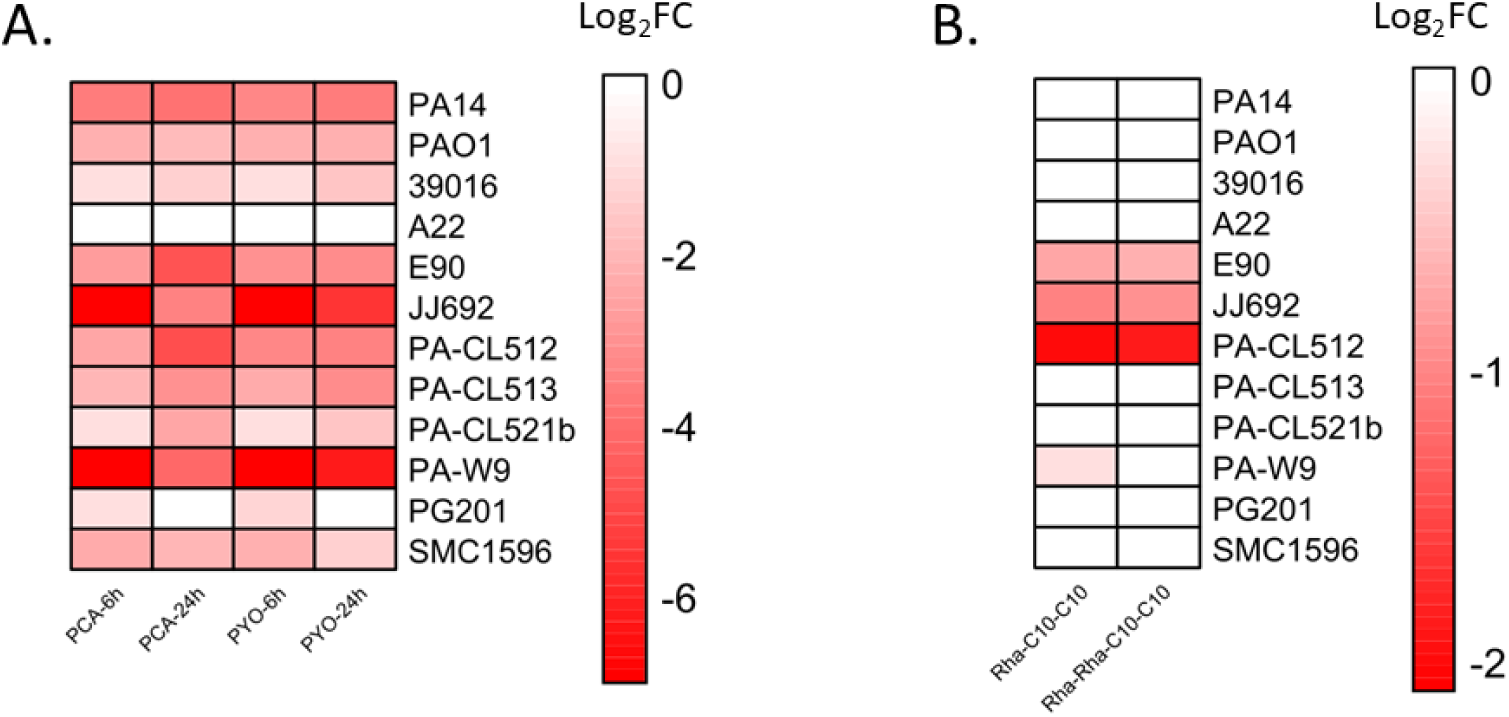
Production of virulence determinants in the Δ*pqsE* mutants relative to their respective WT strain. **(A)** Heatmap of the production of PCA and pyocyanin (PYO) in the Δ*pqsE* mutants relative to their respective WT strain. Concentrations of PCA and pyocyanin (PYO) were measured by LC/MS/MS at 6 and 24 h. (**B)** Heatmap of the production of rhamnolipids in the Δ*pqsE* mutant relative to their respective WT strain. Concentrations of mono-rhamnolipids (Rha-C_10_-C_10_) and di-rhamnolipids (Rha-Rha-C_10_-C_10_) were measured with LC/MS/MS after 24 h of growth in King’s A medium. For both panels, Values are log_2_-transformed ratios (Log_2_ Fold Change (Log_2_FC)) of production in Δ*pqsE* mutants relative to their WT strain (Δ*pqsE*/WT production). Red shades indicate reduced production of metabolites in the mutant, whereas blue indicates increased production. The intensity of the color reflects the magnitude of the change: darker shades indicate greater differences in the logarithmic scale. In panel (A), the color scale ranges from -7 (strong reduction in the mutant) to 7 (strong increase). For visual purposes, the range in panel (B) is from -2 to 2. To enable log transformation and visual consistency, zero values (absence of production) in the mutant were adjusted using a pseudo-count of 1. Ratios were then capped at the lower end of the scale (minimum log_2_ ratio of -7 for panel (A) and -2 for panel (B)). Raw data for these measurements are provided in **Tables S4-S6**.

Since PqsE impacts the activity of RhlR, we chose to look at another RhlR-dependent determinant: rhamnolipid production (11, 56). Rhamnolipids are biosurfactants that play key roles in social motility, biofilm development, and virulence (57). In *P. aeruginosa*, the two principal rhamnolipid congeners are the mono-rhamnolipid Rha-C_10_-C_10_ and the di-rhamnolipid Rha-Rha-C_10_-C_10_ (58). Their biosynthesis is primarily driven by the *rhlAB* operon, which encodes key enzymes involved in the production of these surface-active molecules (56, 59). Given prior evidence suggesting that PqsE plays a role in rhamnolipid production (27, 37, 60) and *rhlAB* transcription through RhlR (24, 34, 61), the production of the two major rhamnolipid congeners in our panel of *pqsE* mutants was investigated using LC/MS/MS quantification.

Rhamnolipid production was reduced in 4 of the 12 *P. aeruginosa* panel strains (**Fig. 2B)**. Additionally, under our culture conditions, loss of *pqsE* in both PAO1 and PA14 did not affect rhamnolipid production, suggesting that any effect of PqsE on the RhlR-mediated *rhlAB* transcription is, at most, very limited. Indeed, previous studies in PA14 have shown only a minimal influence of PqsE on the transcription of the *rhlA* gene, in contrast with an important impact of C_4_-HSL (54). The fact that many *pqsE* mutants do not exhibit reduced rhamnolipid production suggests that the RhlR transcriptional regulator may rely primarily on C_4_-HSL rather than on PqsE for *rhlAB* transcription, as previously suggested (62). In addition, it is important to note that the expression of *rhlAB* is highly dependent on environmental conditions, and various regulatory elements can affect this process (63–65). Therefore, PqsE may be just one of the many factors influencing RhlR-mediated transcription of *rhlAB*.

Curiously, three of the four strains affected by PqsE for rhamnolipid production have a defective LasR protein, and two have a RhlR that functions independently of LasR (**Table 1)** (13, 14). In these strains, we hypothesize that the absence of LasR increases the dependence of RhlR on PqsE for the transcription of the *rhlAB* operon, as RhlR becomes central to the QS hierarchy (29). However, more strains should be studied to verify this hypothesis.

Globally, the role of PqsE in rhamnolipid production varies among strains. While PqsE can influence rhamnolipid production, its effect seems to be strain-specific, indicating that *rhlAB* is not a primary target for PqsE-mediated RhlR activity. Interestingly, no rhamnolipid production defect is observed in the Δ*pqsE* mutants of the PA14 and PAO1 prototypical strains. While previous reports showed limited differences between the WT strain and Δ*pqsE* mutants, these were largely based on the assumption that RhlR-regulated genes are generally also regulated by PqsE. However, more recent data suggest that this is not the case (24, 54). There is also a slight possibility that *rhlAB* does not depend on RhlR in some strains, though this possibility is unlikely, as *rhlA* is considered part of the RhlR core regulon (66). Additionally, while the culture conditions we used are suitable for rhamnolipid production in PA14 and PAO1, we cannot rule out that results could be condition-dependent.

Overall, even if data regarding rhamnolipid production are variable, our results regarding pyocyanin production confirm that PqsE is globally relevant for the regulation of the *rhl* system and the production of virulence determinants, even in LasR-deficient strains (**Table 1**, **Fig. 2)** (29).

### PqsE plays a role in multicellular behaviors of P. aeruginosa strains

RhlR is associated with biofilm formation through the production of various factors, including rhamnolipids and lectins (67, 68). Furthermore, RhlR influences colony biofilm formation (69). Again, since PqsE is linked to RhlR activity, the impact of PqsE on biofilm formation was investigated.

Biofilm formation is a complex process influenced by multiple factors. In *P. aeruginosa*, the biofilm matrix comprises various components, including exopolysaccharides, proteins, and extracellular DNA (67, 70–72). Previous studies based on *in vitro* assays and transcriptomic analyses, primarily conducted with the PAO1 prototypical strain, have shown that PqsE contributes to biofilm formation. Indeed, *pqsE* mutants typically exhibit reduced biofilm production (25, 61). PqsE also influences the structure of colony biofilms in *P. aeruginosa* PA14 (34, 36). Specifically, a *pqsE* mutant exhibits a hyper-rugose phenotype when grown on Congo red agar, primarily due to the loss of phenazine production (34, 73). To further investigate the role of PqsE in this process, we examined biofilm formation in polystyrene plates and on Congo red agar in the strains from our panel and their isogenic Δ*pqsE* mutants.

Our results show variability among the Δ*pqsE* mutants, with some producing more biofilms than their respective WT strains, while others produce less on polystyrene (**Fig. 3)**. Interestingly, under our experimental conditions, the Δ*pqsE* mutant in the PAO1 strain demonstrates increased biofilm formation compared to the WT strain, whereas the opposite effect is seen for the PA14 strain. Overall, an impact of PqsE on biofilm formation *in vitro* appears conserved, with 11 out of 12 isolates showing a significant increase or decrease in biofilm production when *pqsE* is disrupted (**Fig. 3)**.

**Figure 3.**
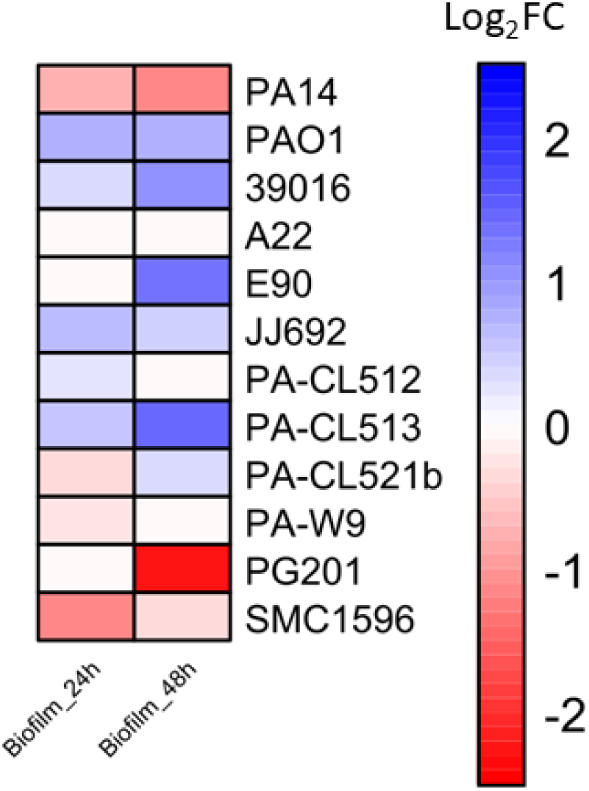
**Heatmap of the production of biofilms in polystyrene plates in the Δ*pqsE* mutant relative to their respective *P****. aeruginosa* WT strain. Biofilm formation (OD_550_) was measured after 24 and 48 hours of incubation at 37°C (74). Values are log2-transformed ratios (Log_2_ Fold Change (Log_2_FC)) of production in Δ*pqsE* mutants relative to their WT strain (Δ*pqsE* /WT production). Red shades indicate reduced biofilm formation in the mutant, whereas blue indicates increased formation. The intensity of the color reflects the magnitude of the change: darker shades indicate greater differences in the logarithmic scale. For visual purposes, the color scale ranges from -2.5 (strong reduction in the mutant) to 2.5 (strong increase). Raw data corresponding to these measurements are provided in **Table S7**.

We also examined colony biofilm formation on Congo red agar. Our results show that PqsE significantly affects the global appearance of colony biofilms in 10 out of 12 strains, even though the resulting morphologies vary rather than consistently displaying the hyper-rugose colony phenotype observed for PA14 (**Fig. 4)**. Strain A22, which does not produce HAQs, shows no difference in biofilm formation on either polystyrene plates or Congo red agar upon *pqsE* deletion, aligning with observations for phenazines and rhamnolipids, in which no differences were noted between the mutant and the WT strain (**Figs. 2 and 3)**.

**Figure 4:**
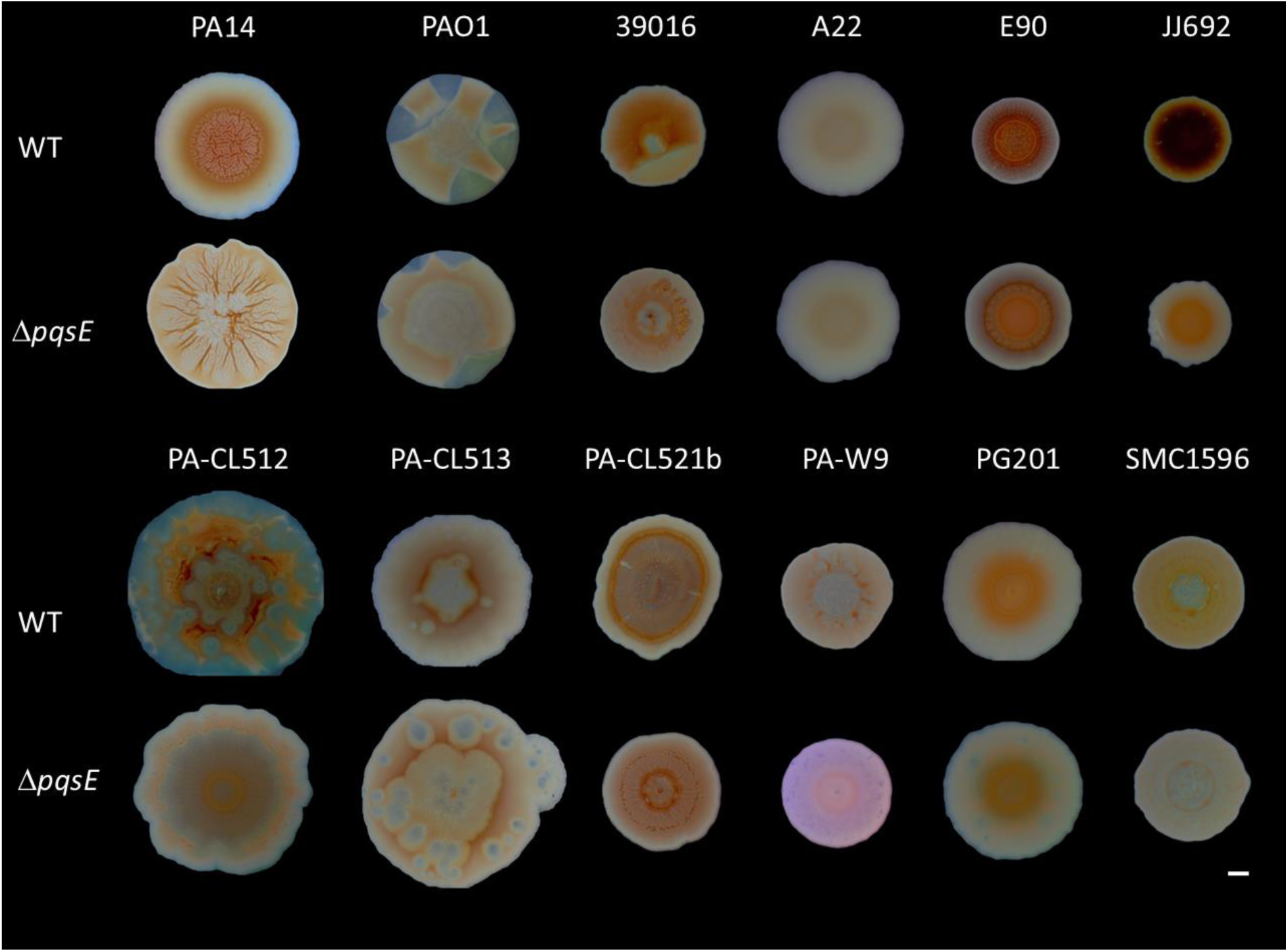
Colony biofilm formation of *P. aeruginosa* strains and their isogenic Δ*pqsE* mutants. Overnight cultures of each strain and their respective Δ*pqsE* mutant were diluted to an initial OD_600_ of 0.05, in TSB medium, and incubated at 37°C with agitation until they reached an OD_600_ of 0.6. 5 μL of each culture were spotted on 1% agar plates containing 1% tryptone, 40 μg/mL Congo Red and 20 μg/mL Coomassie brilliant blue, as described before (75). Pictures were taken with a binocular microscope (Olympus Life Sciences) after 6 days of incubation at room temperature. The scale bar is 2 mm.

The hyper-rugose colony phenotype observed for the PA14Δ*pqsE* mutant has been linked to the absence of phenazine production (73). Since most *pqsE* mutants have reduced PCA and pyocyanin production (**Fig. 2)**, strain-specific differences in phenazine concentrations may influence colony architecture in distinct ways. Indeed, the strains A22 and PG201 show no or very slight effect of PqsE on pyocyanin production and are the only ones that display no significant difference in colony biofilm morphology (**Figs. 2 and 4)**.

The effects of PqsE on biofilm formation vary between strains, likely due to multiple interacting factors. For instance, rhamnolipids play a crucial role in biofilm dispersion and architecture (67, 76–79). In strains where *pqsE* disruption leads to reduced rhamnolipid production, biofilm formation may be enhanced (67, 77). Additionally, recent studies with PAO1 show that PqsE modulates levels of ci-di-GMP through its interaction with the ProE phosphodiesterase (80). A *pqsE* mutant exhibits higher ci-di-GMP levels, which could correlate with enhanced biofilm production (80–82). Conversely, in prototypical strains, PqsE positively regulates genes involved in biofilm formation, including *cupA1*, *lecA* and *lecB* (61). Overall, the interplay of these opposing factors likely contributes to the variations in biofilm formation across different strains, highlighting the complexity of this process.

We also examined swarming motility, another multicellular behavior depending on RhlR. Swarming motility is a collective behavior characterized by the rapid and coordinated movement of groups of cells on a semi-solid surface (83–86). To swarm, *P. aeruginosa* requires both rhamnolipid production and a functional flagellum (87, 88). PqsE is thought to be involved in this social behavior, as a *pqsE* mutant in the PAO1 prototypical strain shows reduced swarming motility under some conditions (25). Since swarming motility requires rhamnolipids, we assessed swarming motility in the four strains where the Δ*pqsE* mutation led to reduced rhamnolipid production (E90, JJ692, PA-CL512 and PA-W9) (**Fig. 2B)**. We compared the swarming patterns of the WT strains and their Δ*pqsE* mutants to that of the PA14 strain, for which rhamnolipid production remains unaffected by *pqsE* depletion. Among these strains, three (E90, JJ692 and PA-CL512) exhibited varying degrees of swarming motility reduction of their Δ*pqsE* mutants (**Fig. 5)**, consistent with our rhamnolipid quantification results (**Fig. 2B)**. The remaining strain (PA-W9) displayed a minor alteration in the swarming pattern but no significant change in swarm coverage (**Fig. 5)**. Indeed, this strain only showed a very slight decrease in mono-rhamnolipid (Rha-C_10_-C_10_) production in the Δ*pqsE* mutant, which may not be sufficient to have a significant impact on the swarming behavior (**Fig. 2B)**. Our results confirm that PqsE can indeed be involved in swarming motility, most likely through its impact on rhamnolipid production.

**Figure 5:**
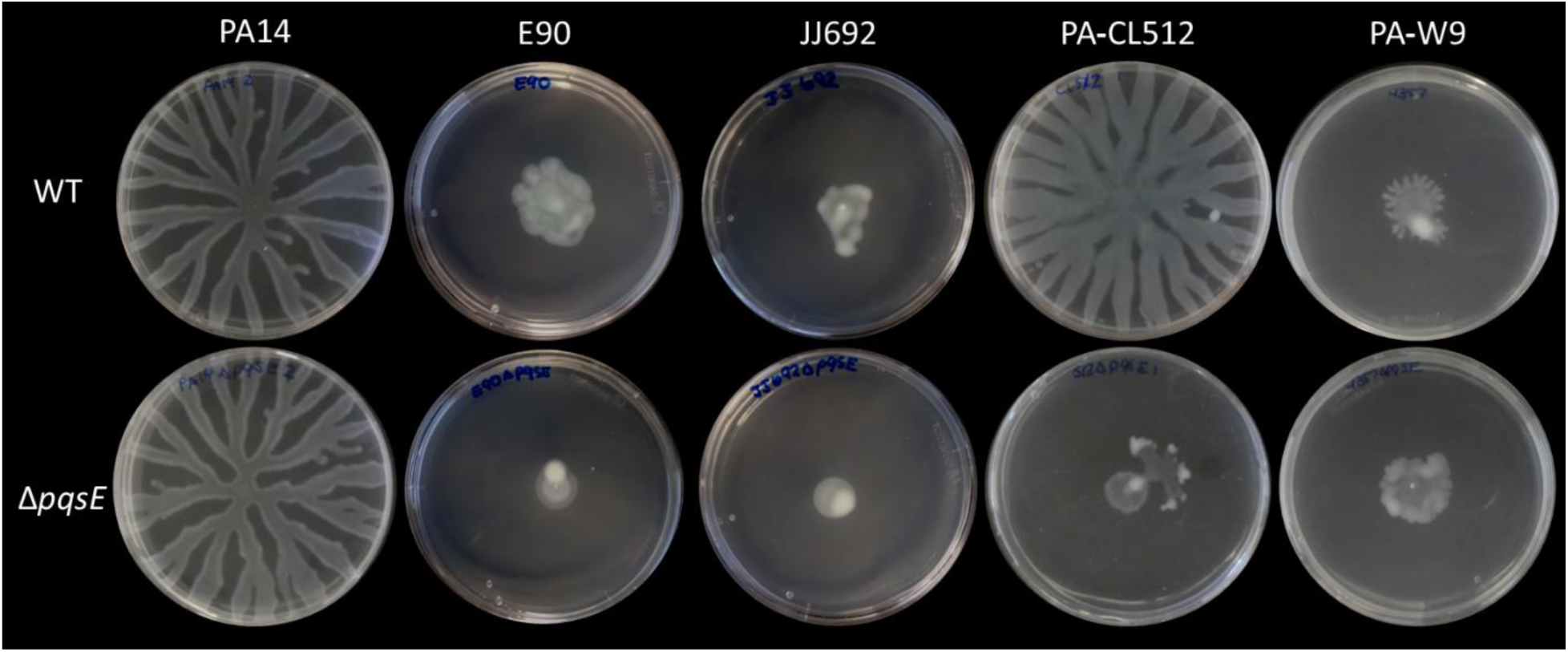
Swarming motility of four *P. aeruginosa* isolates exhibiting reduced rhamnolipid production in their Δ*pqsE* mutant. PA14 and its isogenic Δ*pqsE* mutant were used as negative controls. Images were acquired after 24 hours of incubation at 37°C on semi-solid MD9CAA medium.

Overall, PqsE can play a significant role in RhlR-dependent multicellular behaviors, including swarming motility and biofilm development. Given the complexity of biofilm regulation, PqsE likely influences multiple regulatory pathways, leading to strain-specific outcomes. Despite the variability of phenotypes between strains, PqsE remains a key factor in shaping multicellular structures in *P. aeruginosa* isolates.

### The function of PqsE in HAQ-defective strains remains unknown

After confirming the genetic conservation of *pqsE*, we assumed it was transcribed in all our strains. Among the 12 *P. aeruginosa* isolates from our panel, 11 produce HAQs (13, 14) (**Table 1)**. In those strains, there is likely transcription of the *pqs* operon and, consequently, *pqsE* (18). Supporting this, all *pqsE* mutants of HAQ-producing strains exhibit differences in at least one of the tested phenotypes compared to their respective WT strains (**Fig. 2**, **Fig. 3**, **Fig. 4)**. Strain A22 does not produce HAQs, but still produces pyocyanin (14) **(Table S5)**. Notably, previous studies have shown that *pqsE* can still be expressed in some strains that lack HAQ production yet still produce pyocyanin (62, 89). However, A22 is the only isolate from our panel with no significant differences with the Δ*pqsE* mutant in any tested virulence determinants, including the production of PCA, pyocyanin, and rhamnolipids (**Fig. 2)**. The same goes for multicellular behaviors (**Fig. 3**, **Fig. 4)**. This raised the possibility that *pqsE* may not be transcribed in this strain under our experimental conditions, and these factors would be produced without an involvement for a PqsE homologue. To investigate this further, we performed RT-PCR on *pqsE* in the A22 strain. The PA14 prototypical strain as well as PA14 carrying a non-polar mutation in *pqsA* were used as controls. This mutant can still express *pqsE* and produce detectable pyocyanin while lacking HAQ production, which represents an ideal control for the A22 HAQ-defective strain (**Fig. 6, data not shown)**. Our results confirm that A22 does transcribe *pqsE*, reinforcing that it can occur independently of HAQ production, as previously reported for some *P. paraeruginosa* strains (**Fig. 6)** (62, 89). However, these results suggest that in some HAQ-negative strains, PqsE-dependent phenotypes, such as pyocyanin production, may be completely independent of PqsE. Previous studies have reported only partial dependence of RhlR towards PqsE, supporting the possibility of a subset of strains having PqsE-independent RhlR activity (62). Globally, the role of PqsE, as well as the whole QS regulatory circuitry, remains largely unknown in HAQ-defective strains. Given that HAQ-negative strains represent approximately 20% of all *P. aeruginosa* isolates (13, 14), the mechanisms underlying this adaptation warrant further investigation.

**Figure 6.**
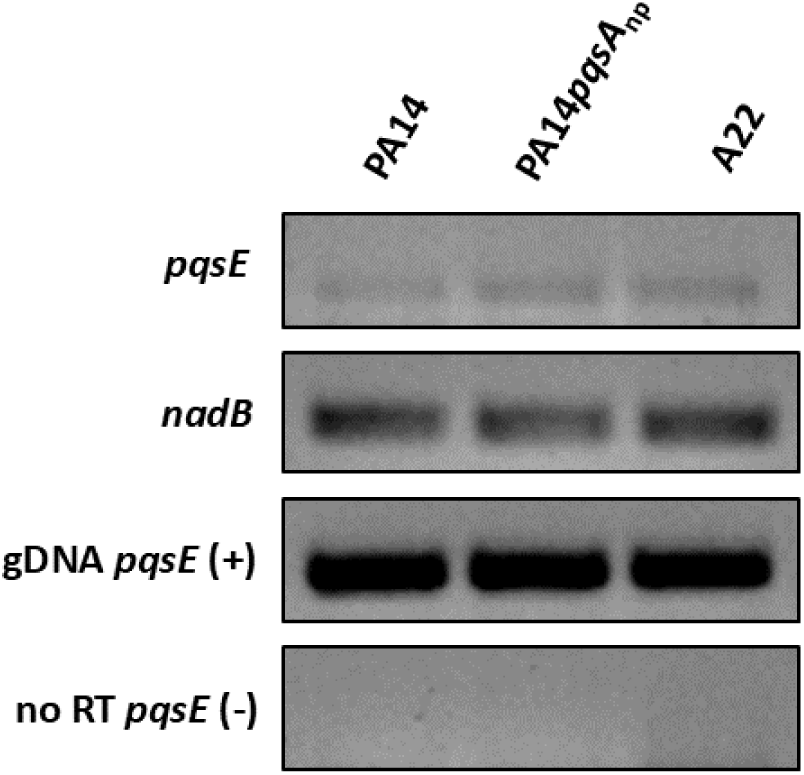
: RT-PCR of *pqsE* transcription in A22 strain. A 2% agarose gel displaying RT-PCR amplification of *pqsE* mRNA levels from strain A22 alongside the reference strain PA14, and a PA14*pqsA* nonpolar mutant (PA14*pqsA*_np_). Cultures were grown in King’s A medium at 37°C with agitation for 6h. The *nadB* gene was used as a housekeeping control. Genomic DNA (gDNA) served as a positive control (+), while a no-reverse transcriptase (no-RT) sample was included as a negative control (-).

### HAQ production can also be a PqsE-dependent determinant

In addition to regulating the production of virulence factors through RhlR, PqsE has a thioesterase activity. PqsE catalyzes the conversion of 2-aminobenzoylacetyl-coenzyme A (2-ABA-CoA) to 2-aminobenzoylacetate (2-ABA), which is ultimately used to produce HHQ and PQS, the key ligands of the transcriptional regulator of the PQS system, MvfR (16, 18, 23). However, some studies have shown that PqsE is not strictly essential for this function, as a *pqsE* mutant in PA14 and PAO1 are reported to produce WT levels of HHQ and PQS (18, 19, 23). This was explained by the fact that other thioesterase enzymes, such as TesB, can compensate for PqsE’s enzymatic activity (23). To verify whether what was observed in prototypical strains remains true in other strains, we assessed HAQ levels in our panel (**Fig. 7)**. Specifically, we quantified HHQ and PQS production at two distinct time points in HAQ-producing strains (i.e. A22 was not included). These time points were selected based on prior studies analysing HAQs in large collections of diverse strains (13, 14).

**Figure 7:**
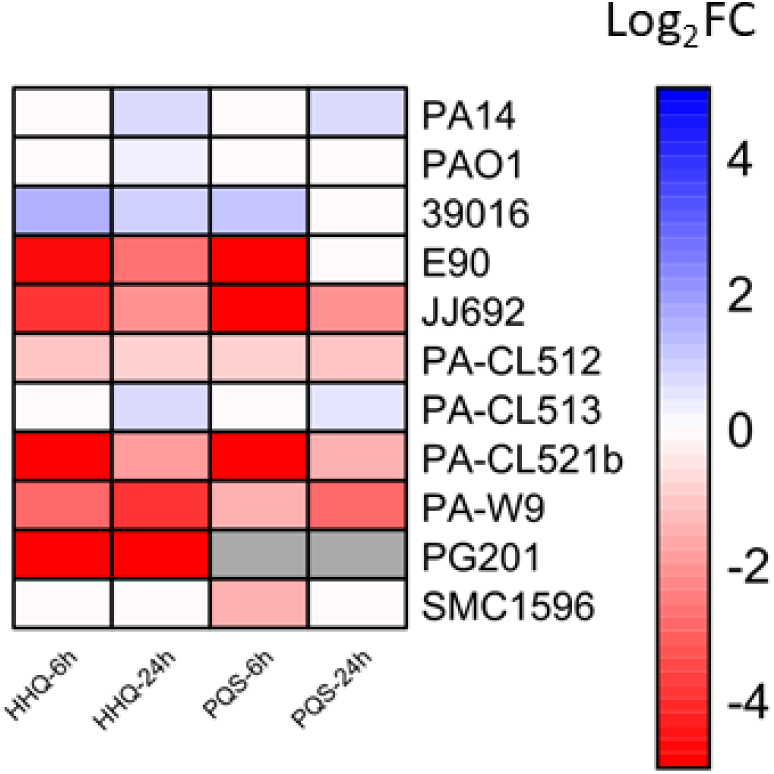
Heatmap of the production of HHQ and PQS in the Δ*pqsE* mutants compared to the *P. aeruginosa* WT strains. Concentrations of HAQs (HHQ and PQS) were measured by LC/MS/MS at 6 and 24 h. Values are log_2_-transformed ratios (Log_2_ Fold Change (Log_2_FC)) of production in Δ*pqsE* mutants relative to their WT strain (Δ*pqsE*/WT production). Red shades indicate reduced production of metabolites in the mutant, whereas blue indicates increased production. Gray indicates no production. The intensity of the color reflects the magnitude of the change: darker shades indicate greater differences in the logarithmic scale. The color scale ranges from -5 (strong reduction in the mutant) to 5 (strong increase). To enable log transformation and visual consistency, zero values (absence of production) in the mutants were adjusted using a pseudo-count of 1. Ratios were then capped at the lower end of the scale (minimum log_2_ ratio of -5). Raw data for these measurements are provided in **Tables S8-S9**.

Surprisingly, most strains have significantly lower HHQ and/or PQS production in Δ*pqsE* mutants compared to their respective WT strains. In some cases, HAQ production was completely abolished in the absence of *pqsE* (**Fig. 7)**. This suggests that other thioesterase enzymes may not be as effective in most strains to fully compensate for the loss of PqsE activity, in contrast with the previous suggestion (23), and that this corresponds to an expected involvement for the presence of *pqsE* in the HAQ biosynthetic gene cluster.

Also unexpectedly, a subset of strains (39016, PA-CL513, PAO1 and PA14) behaved similarly: the PqsE enzymatic activity did not appear to be essential for HAQ production at either time point. Actually, in these strains, HAQ levels were even higher in the mutants than in the parental strain, with an accumulation especially observed in the later stages of growth (24h) (**Fig. 7)**. Interestingly, aside from PAO1, the isolates that behaved similarly in absence of *pqsE* in terms of HAQ production were the ones clustering with PA14 in our core genome phylogenetic analysis (**Fig. 1, Fig. S4)**. This suggests that these strains may share a common genetic evolution, either the loss of a gene or the presence of an additional gene, which compensates for the thioesterase enzymatic activity of PqsE. Furthermore, PqsE exerts a negative autoregulatory feedback loop on the transcription of the *pqsABCDE* operon in prototypical strains again through its impact on RhlR (25). However, whether this is also the case in other *P. aeruginosa* strains remains to be determined.

These findings highlight the crucial role of PqsE thioesterase activity in the synthesis of HAQs across many *P. aeruginosa* strains, contradicting previous conclusions that were based on limited assessment in prototypical strains. Notably, this challenges the widely accepted notion that PqsE is not involved in HAQ production.

## CONCLUSION

The thioesterase PqsE moonlighting as a quorum sensing effector protein has been extensively studied in recent years, particularly concerning its role in promoting the RhlR regulon through a protein-protein interaction (24, 26, 29–31). Because of its impact on virulence and its presence restricted to very few species such as *P. aeruginosa*, PqsE could be an interesting target for the development of specific anti-virulence therapies. To reinforce its appeal, it is important to show that the presence and functionality of PqsE is a conserved trait among *P. aeruginosa* strains outside of the few prototypical strains described in the literature. In this study, we explored the conservation and function of PqsE in a panel of 12 genetically diverse environmental and clinical strains of *P. aeruginosa,* including PA14 and PAO1. To achieve this, we looked at the conservation of the *pqsE* gene as well as the impact on phenotypes considered to PqsE-dependent.

Our findings confirm the genetic conservation of *pqsE* and *rhlR*, consistent with previous reports (46, 50). Indeed, the few identified SNPs do not affect the amino acid sequence. Furthermore, our phenotypical survey of the *pqsE* mutants from our panel reveals that while the impacts might vary, phenotypes affected by PqsE remain consistent across strains.

Since our strain panel is genetically diverse and represents variations within QS systems (14, 29), our results likely reflect the diversity of the PqsE function, emphasizing the importance of studying multiple isolates before reaching general conclusions. The reasons why, in some contexts, a subset of strains do not require PqsE to activate the RhlR regulon and other known PqsE targets remain unclear and warrant further investigation. The possibility that some strains retain full RhlR activity independently of PqsE remains of interest.

The interaction between PqsE and RhlR has been proposed as an interesting target for anti-virulence therapies due to its conservation and functional importance in producing some virulence factors in prototypical strains (26, 30, 31). Further studies on additional PqsE targets are needed to gain a better understanding of the role of this protein in QS regulation. Moreover, testing environmental and clinical *P. aeruginosa* strains and their isogenic Δ*pqsE* mutants in animal models will provide deeper insights into the role of PqsE in virulence and confirm its potential as a therapeutic target.

## MATERIAL AND METHODS

### Strains and growth conditions

The isolates selected for this study were collected from the environment and various chronic and acute clinical infections. Clinical strains were obtained from the International *Pseudomonas* Consortium Database (IPCD) (38), while environmental strains were previously isolated from hospital sinks and soils. Prototypical strains UBCPP-PA14 (90) and PAO1-N (University of Nottingham collection) were also included **(Table S1)**. A detailed list of the selected isolates and some of their genetic and phenotypic characteristics is presented in **Table 1**. Unless indicated otherwise, Tryptic Soy Broth (TSB) medium (BD Difco) was used for the routine growth of bacteria, and cultures were incubated at 37°C in a TC-7 roller drum (New Brunswick) at 150 r.p.m.

### Whole genome sequencing

For each *P. aeruginosa* strain of this study, DNA was extracted as previously described in Freschi et al. (2015), as part of the International *Pseudomonas* Consortium Database project (38). Briefly, DNA extraction was performed with the DNeasy Blood and Tissue Kit (QIAGEN) following the manufacturer’s recommended protocol for Gram-negative bacteria. The genomic DNA was subsequently fragmented into 450 ± 70 bp inserts with a Covaris M220 ultrasonicator. These libraries were then barcoded with TruSeq adapters and sequenced on an Illumina MiSeq apparatus in 2 x 300 bp mode.

### Genome assembly, annotation, and sequence analyses

Illumina MiSeq reads were assembled with the A5-miseq pipeline v2015.05.22 (91). After assembly, genomes were annotated with Prokka v1.14.6 (92) with default parameters except for the use of a Prokka genus-specific database (--usegenus --genus Pseudomonas). Subsequently, MAFFT v7.511 was used to align the *pqsE* and *rhlR* sequences and identify mutations (93). Alignments were then visualized with Jalview v2.11 (94).

### Generation of phylogenetic tree and average nucleotide identity (ANI) heatmap

A core gene alignment was done with PyMLST v2.1.6 (95) and the cgMLST.org core genome MLST scheme for *P. aeruginosa* (https://www.cgmlst.org/ncs/1000/schema/Paeruginosa2033/). Monomorphic and underrepresented sites were filtered from the nucleotide matrix with BMGE v1.12 (96). Then, a maximum likelihood phylogenetic tree was constructed using IQ-TREE v2.3.6 (97), with statistical support from 1,000 bootstrap replicates. The optimal substitution model was automatically inferred by IQ-TREE (GTR+F+I+G4). The resulting tree was visualized with FigTree v1.4.4 (http://tree.bio.ed.ac.uk/software/figtree/). ANI calculations were done with pyani v2.1.6 (https://github.com/widdowquinn/pyani) using the “ANIm” all-versus-all comparison mode, with default parameters. The resulting percentage matrix was aligned with the nodes of the phylogenetic tree described above.

### Plasmid construction

The pMT01 plasmid (pEX18Gm-Δ*pqsE*) was constructed by modifying the existing *sacB*-containing suicide plasmid pEX18Ap-Δ*pqsE*, which contains a 570-bp deletion allele of the *pqsE* gene (18). The pEX18Ap-Δ*pqsE* plasmid was digested with *EcoRI* and *BamHI*, releasing a 1905-bp Δ*pqsE* fragment. This fragment was purified and ligated into the pEX18Gm backbone with the same restriction enzymes (98), using the T4 DNA ligase (NEB). A comprehensive list of the plasmids used in this study is provided in **Table S10**.

### Construction of in-frame deletion mutants

An allelic replacement method adapted from Hmelo et al. (2015) was used to create deletion mutants in the *pqsE* gene (99). Suicide vector pMT01 was introduced into recipient *P. aeruginosa* strains by conjugation with donor auxotrophic *Escherichia coli* strain χ7213 on plates containing 50 µg/mL diaminopimelic acid (DAP). Merodiploid cells were selected on media containing gentamicin at pre-determined concentrations for each strain. Double crossover mutants were isolated through sucrose counterselection, and the mutation in *pqsE* was confirmed by PCR, using primers listed in **Table S11**.

### Quantification of secondary metabolites and quorum sensing molecules

For the quantification of PCA, pyocyanin, and HAQ molecules (HHQ and PQS), overnight cultures of wild-type (WT) strains and their isogenic Δ*pqsE* mutants were diluted to an OD_600_ of 0.05 in King’s A broth supplemented with 100 µM of FeCl_3_ (100). Cultures were prepared in triplicates and incubated at 37°C under agitation for 6 and 24 hours, with sampling time points selected based on previous studies (13). To extract metabolites, 375 µL of acetonitrile containing tetradeuterated 4-hydroxy-2-heptylquinoline (HHQ-d4) as an internal standard was added to 1.5 mL of culture sample. The suspension was vortexed and centrifuged for 10 minutes at 17,000 x *g* to pellet the bacteria. Supernatants were transferred into vials and analyzed using a liquid chromatography/mass spectrometry/mass spectrometry (LC/MS/MS) method, as described previously (101).

For rhamnolipid quantification, a similar procedure was followed for the preparation of samples with some modifications. Only the 24h time point was used to measure rhamnolipid accumulation, and supernatants were diluted 20-fold before analysis. The quantification of rhamnolipids in the supernatant was performed by LC/MS/MS, as described before (57).

Due to the tendency of many clinical and environmental strains to form clumps, precluding the use of absorbance to assess growth, relative concentrations of secondary metabolites and HAQ molecules were normalized to the total protein content of the cell pellet collected from the whole culture at the time of sampling. The pellets were resuspended in 0.1 N NaOH and incubated at 70°C for 1h. Total protein concentrations were measured using the Bradford protein assay (Bio-Rad, Montreal, Canada), with bovine serum albumin (BSA) serving as a standard.

### Reverse transcription polymerase chain reaction (RT-PCR)

Overnight cultures were diluted in triplicates to an OD_600_ of 0.05 in King’s A medium and incubated at 37°C with agitation. Cells were harvested after 6 hours of growth. Total RNA was extracted using Aurum™ Total RNA Mini Kit (Bio-Rad Laboratories). To eliminate any residual DNA, the extracted RNA was treated with TURBO DNA-Free Kit (Ambion, Life Technologies). Reverse transcription was performed using iScript™ gDNA Clear cDNA Synthesis Kit (Bio-Rad Laboratories). A portion of the *pqsE* gene was amplified using specific primers **(Table S11)**, and the resulting PCR products were analyzed by electrophoresis on a 2% agarose gel. The *nadB* gene was used as a housekeeping control (102), and no-RT and gDNA controls were also included.

### Swarming motility

Swarming motility assays were performed as previously described (103). Briefly, 20 mL of M9DCAA medium with 0.5% Bacto-agar (Difco) was poured into 100 mm Petri dishes and allowed to dry in a laminar flow cabinet. Overnight cultures were adjusted to an OD_600_ of 3.0, and 5 μL was inoculated in the center of an agar plate. Plates were then incubated at 37°C for 16 hours. Each strain was tested in triplicates.

### Biofilm formation

Biofilm formation was quantified using crystal violet staining, as described before (74). Briefly, overnight cultures of all strains and their isogenic Δ*pqsE* mutants were diluted to an OD_600_ of 0.05 in M63 minimal medium supplemented with 1 mM magnesium sulfate, 0.2% dextrose, and 0.5% casamino acids. For each strain, a 100 μL aliquot was added to five wells of polystyrene 96-well plates, which were incubated at 37°C for 24 and 48 hours. After incubation, plates were rinsed thoroughly with water, and 125 μL of 0.1% crystal violet was added to each well. Following a 15-minute incubation at room temperature, the plates were rinsed again, and the bound dye was solubilized in 125 μL of 30% acetic acid. Absorbance was measured at 550 nm using a Cytation microplate reader (Biotek).

### Colony biofilm formation

Overnight cultures of WT strains and Δ*pqsE* mutants were diluted to an OD_600_ of 0.05 in TSB medium and incubated at 37°C under agitation until they reached an OD_600_ of 0.6. As described before, 5 uL of each dilution was inoculated onto 1% agar plates containing 1% tryptone, 40 μg/mL Congo Red, and 20 μg/mL Coomassie brilliant blue (75). Plates were incubated at room temperature for 6 days. Pictures of colony biofilms were taken with a binocular microscope (Olympus Life Science). This experiment was repeated at least twice for each strain.

### Statistical analyses

Statistical significance in metabolite production or gene expression between the WT strains and their respective Δ*pqsE* mutants was determined using the Holm-Sidak method, with a significance threshold (*p-value*) of 0.05 (GraphPad Prism). When a significant difference was detected, the production or expression levels in the Δ*pqsE* mutants were divided by those of the WT strains to calculate the proportion. In cases where no significant difference was observed, the production of the Δ*pqsE* mutant relative to the WT strain was assumed to be 1. Proportions were subsequently log2-transformed for analysis. A small pseudocount of 1 was added for instances where the mutant’s production or expression was zero to enable log transformation. Log2-transformed proportions were capped at a subjective minimum threshold (indicating the absence of production) to prevent extreme values. Heatmaps were generated with the R software (104), using the package ‘’pheatmap’’ (105).

## Acknowledgements

Many thanks to Ajai A. Dandekar (University of Washington) for supplying strain E90 and Giordano Rampioni (Università degli Studi Roma Tre) for strain PAO1-N and its isogenic Δ*pqsE* mutant. The authors also thank Sandrine Gervais and Maude Dagenais Roy for their technical support. This work was funded by Canadian Institutes of Health Research (CIHR) operating grants MOP-142466, 482990 and 508306. MCT was recipient of a Canada Graduate Scholarship – Doctoral program from the Natural Sciences and Engineering Research Council of Canada (NSERC).

